# Concept2Brain: An AI model for predicting subject-level neurophysiological responses to text and pictures

**DOI:** 10.1101/2025.08.04.668476

**Authors:** Alejandro Santos-Mayo, Faith Gilbert, Arash Mirifar, Anna-Lena Tebbe, Ruogu Fang, Mingzhou Ding, Andreas Keil

**Affiliations:** Laboratory of Brain, Body, and Behavior, University of Florida, Gainesville, FL, USA; Department of Psychology, University of Florida, Gainesville, FL, USA; Center for Cognitive and Computational Neuroscience, Complutense University of Madrid, Madrid, Spain; J. Crayton Pruitt Family Department of Biomedical Engineering, University of Florida, Gainesville, FL, USA

## Abstract

The current growth of artificial intelligence (AI) tools provides an unprecedented opportunity to extract deeper insights from neurophysiological data while also enabling the reproduction and prediction of brain responses to a wide range of events and situations. Here, we introduce the Concept2Brain model, a deep network architecture designed to generate synthetic electrophysiological responses to semantic/emotional information conveyed through pictures or text. Leveraging AI solutions like CLIP from OpenAI, the model generates a representation of pictorial or language input and maps it into an electrophysiological latent space. We demonstrate that this openly available resource generates synthetic neural responses that closely resemble those observed in studies of naturalistic scene perception. The Concept2Brain model is provided as a web service tool for creating open and reproducible EEG datasets, allowing users to predict brain responses to any semantic concept or picture. Beyond its applied functionality, it also paves the way for AI-driven modeling of brain activity, offering new possibilities for studying how the brain represents the world.

## 1. Introduction

One of the most significant scientific milestones of the 1920s was when Hans Berger recorded the first human scalp electroencephalogram (EEG) in 1924, laying the foundation for understanding the neurophysiological dynamics of the human brain^1^. Initially confined to medical and clinical research, EEG has since evolved into a versatile tool with applications extending beyond neuroscience into fields such as economics, marketing^2^, rehabilitation^3^, and human-computer interaction^4^, including brain-computer interface (BCI) technology^5^. More recently, numerous artificial intelligence (AI)-driven approaches to EEG data analysis have emerged ^6–8^. These methods are widely employed in research laboratories to classify EEG signals, to enhance real-time brain decoding^6,9–11^, and to optimize BCI performance^12,13^. This synergy between EEG and AI reflects a broader trend, with recent years showing an exponential rise in AI-driven solutions, applied to both scientific research and real-world problems.

While earlier AI approaches in neuroscience focused on analyzing brain activity for data classification and computational modeling, a more recent and promising direction involves creating artificial systems that can generate or reproduce brain-like data^12–14^. Instead of decoding the brain activity to classify or reconstruct a given stimulus, event, or condition, this approach intends to encode the stimulus to generate bio-realistic brain activity. Well-validated encoding models open new possibilities to understand neural processes by enabling researchers to test hypotheses in simulated environments. However, despite its potential, encoding models for EEG remain relatively underexplored, with many questions regarding their ability to accurately replicate the complexity of real brain activity.

To address this problem, the present report introduces the Concept2Brain model, a deep neural network architecture designed to generate synthetic EEG signals from either images or text. Specifically, the model has been trained to reconstruct visual event-related brain potentials (ERPs) associated with the perception of semantically rich scenes varying in content. ERPs are voltage changes caused by electrophysiological processes in the brain, recorded from the scalp. They are reflective of activity at several hundreds of thousands of neurons in the outer layer of the brain, the so-called cerebral cortex. This activity is quantified relative to specific sensory or behavioral events by repeating those events many times and averaging the resulting voltage time series. Providing millisecond temporal resolution, ERPs present typical time courses or neuroelectric activity that reflect the brain’s response to specific events, evident in positive and negative voltage deflections^17,18,19,20,21–23^. These electrophysiological elements have proven consistent and reliable across animal and human studies. Specifically, the topography and amplitude of late ERP dynamics has been linked to the brain’s processing of semantic concepts, including those varying in emotional content^21,22,24,25^ (see Figure 2, right column for an example). Thus, brain dynamics as captured by ERPs have the potential to elucidate a given observer’s cognitive and affective processing of real-world concepts. If information is systematically shared between real world concepts expressed in media and text and their representation in the human brain, then it should be possible to translate semantic content from the domain of graphical or language representations into the realm of brain activity.

The Concept2Brain model is designed to perform this function, using ERP data. It is available as an open cloud platform (http://concept2brain.brainbodybehavior.science) where brain responses for different synthetic human participants can be generated based on user-entered prompts, which can be written text or digital images. The generation of synthetic brain data holds transformative potential across diverse fields. By generating large, diverse, and customizable datasets, this technology overcomes temporal and financial barriers associated with costly and difficult EEG/ERP data acquisition. It thus enables researchers with a wide spectrum of backgrounds and interests to generate simulated data, facilitating investigating novel hypotheses and conducting simulation studies as well as power analyses. The present report details the generation and validation of this novel resource and discusses its many potential applications.

## 2. Methods

The Concept2Brain model consists of a deep neural network architecture trained to generate simulated electrophysiological signals in response to a wide range of semantic and emotional content. The model has been trained using EEG data from human observers, viewing several hundreds of naturalistic scenes varying in semantic and emotional content. In the following sections, we give a detailed description of the deep learning architecture, its training, and its performance evaluation.

### 2.1. Concept2Brain: a Cross-Domain Conditioned Variational Autoencoder model

The Concept2Brain model comprises a cross-domain conditioned variational autoencoder architecture based on deep learning networks. This architecture can be divided into two main components, performing three tasks: (1) The first component is a conditioned variational autoencoder (cVAE)^15–17^, which is a neural network designed to extract latent dimensions from the electrophysiological data and reconstruct the original data from the resulting low dimensional latent space. (2) The second component is a cross-domain deep network, which is a dense multi-layer network translating information contained in the conceptual domain to the electrophysiological domain. This network converts pictures or text elements captured by an AI solution into a single vector embedding and then translates this concept-level latent information into the latent electrophysiological space of the cVAE. (3) Finally, both deep learning architectures are connected in the Concept2Brain model, converting picture or text information into simulated EEG recordings. In the following, we outline the properties of the components of our model.

#### 2.1.1. The Conditioned Variational Autoencoder model for compressing and predicting electrophysiological data

The cVAE is a type of autoencoder (AE)^18–20^, a deep neural network in which the output data mirror the input data as closely as possible. Importantly, in the center of the AE architecture, hidden network layers converge into a smaller layer called the “bottleneck”. This low-dimensional layer is also referred to as the “latent space” and represents a compressed version of the data that contains a sufficient amount of information needed for the reconstruction of the input data (Figure 1.1, light purple vector). In the present cVAE, both input and output data consist of spatio-temporal matrices of ERP signals having a size of 129 (recording channels) by 750 (time points, representing 1.5 s at the sampling frequency of 500 Hz). These data are passed through an encoder, bottleneck, and decoder. Both the encoder and decoder architectures have three convolutional layers^21,22^, designed to extract spatial and temporal features from input data. In EEG and ERP analysis, convolutional layers are particularly useful for capturing the temporal dynamics of brain signals^23,24^. The convolutional layers of the Concept2Brain model extract the electrophysiological latent space, effectively compressing the spatio-temporal information of the EEG signal.

**Figure 1.**
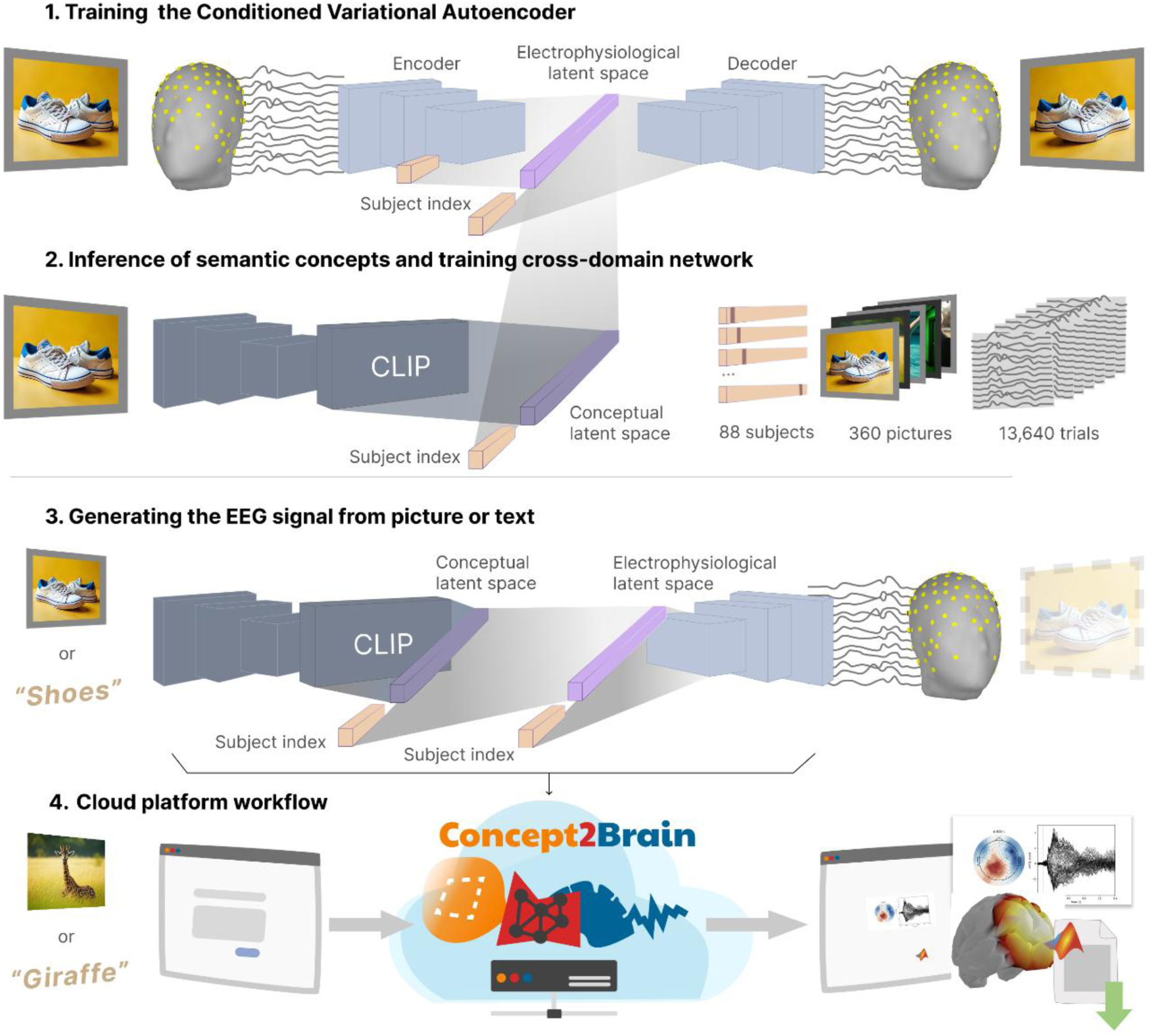
Development and deployment scheme of the Concept2Brain model. 1) Training phase of the cVAE architecture for extracting the electrophysiological latent space from the EEG datasets. The blue boxes, light purple vector and light orange vectors refer to the convolutional neural layers, the bottleneck of the network and the vector of subject ID, respectively. 2) The CLIP model by OpenAI is used to parse the semantic content of pictures into a multidimensional conceptual latent space. Then, a neural network is trained as a cross-domain model, linking the conceptual and electrophysiological latent spaces. 3) The Concept2Brain model leverages the cross-domain cVAE models to translate pictures or text into synthetic EEG signals. 4) This model is deployed in a cloud-computing platform for open-access use of the model without any hardware nor software requirements.

Unlike a standard AE, which learns a low-dimensional latent representation for input data but lacks a well-structured latent space for meaningful sampling, the cVAE model enables the synthesis of new data samples, the cVAE model enables the synthesis of new data samples^16^. Specifically, a standard AE encodes data into a bottleneck layer and reconstructs it without enforcing any distributional constraints on the latent space. As a result, the latent space may be sparsely populated and discontinuous, making it difficult to interpolate or sample new data points that resemble the training distribution—in the present case EEG data. In contrast, the variational AE (VAE), explicitly models the latent space as a structured distribution (typically Gaussian) conditioned on auxiliary information, ensuring that samples drawn from the latent space can be decoded into realistic and semantically consistent data— potentially reflecting biologically plausible features in domain-specific applications^16,25^. The Variational Autoencoder (VAE) addresses this limitation by introducing a probabilistic framework that regularizes the latent space to follow a standard Gaussian distribution. In the VAE architecture, the bottleneck contains two parallel layers that output the mean (*μ*) and standard deviation (*σ*) of the latent variable distribution. During training, a latent vector *z* is sampled using the reparameterization trick: *z* = *μ* + *σ* ⋅ *ϵ z* where *ϵ* ∼ *N*(0, *I*). This approach enables differentiable sampling and ensures that points drawn from the latent space correspond to realistic reconstructions, thereby facilitating meaningful data generation. As a result, the latent dimension becomes interpretable, because it describes features in the electrophysiological space. Here, we used a size of 256 units for the bottleneck (both μ and σ layers). This value was chosen to enable the representation of the variability across participants in the empirical data used for training. As a result, the VAE architecture enables the generation of new data by introducing 256 z-score values at the beginning of the decoder component of the VAE network and forwarding them through the architecture, until the output layer (see Figure 1.1).

The architecture also represents variability across participants. To this end, the cVAE incorporates additional information (conditioning variables) immediately before and after the bottleneck (Figure 1.1, light orange)^17^. In our architecture, the conditioned information represents a subject to whom the EEG input belongs. Thus, a vector of *n* x 1 (in our example, the total *n* is 88 participants) is added with a value of 1 in the index of the subject (one-hot-vector) and all other values set to zeros. This vector is added to the model just before and after the bottleneck layers (i.e. the latent vectors). By including this information, the cVAE forces the bottleneck to learn general electrophysiological properties by compressing the EEG signals regardless of the idiosyncratic properties specifically related to each subject. Then, the conditioning of the cVAE model allows the generation of new synthetic EEG recordings for each of the 88 subjects reproducing some of their idiosyncratic electrophysiological responses. Thus, it is possible to effectively create simulated multi-participant studies, illustrating how different brains would respond to the input picture or text provided.

#### 2.1.2. The Cross-Domain bridge between the concept-level and electrophysiological latent spaces

The Concept2Brain model provides a link between electrophysiological information derived from cortical activity and concept-level content. To this end, the model relies on a third-party AI solution for extracting the conceptual representation of real-world stimuli such as pictures, texts, or audio. Currently, AI solutions and alternatives are rapidly evolving and increasing. Crucially, the Concept2Brain idea does not rely solely on one specific model, and its future versions may adopt different models as they advance or offer improved features related to diverse objectives.

One of these AI solutions is the Contrastive Language-Image Pre-Training (CLIP) model^26^ (Figure 1.2). We selected this model developed by OpenAI (accessible from https://github.com/openai/CLIP) because it captures the representations of images^27^ and texts^28^ from large-scale datasets and embeds them in a common latent space. Specifically, the CLIP model compresses the information contained in the input (pictures or text) into a multimodal vector of 512 dimensions. In Concept2Brain, this representation is called the conceptual latent space.

To connect both the conceptual representational space, obtained from CLIP, and the electrophysiological representational space extracted using our EEG-cVAE architecture, we applied a cross-domain multi-layer neural network^29,30^ (Figure 1.2, shaded network between top and middle rows). This part of the architecture consists of three dense fully connected layers of neurons. It abstracts non-linear information from the 512-size CLIP output and estimates the 256-size latent vector of the cVAE, thus mapping the conceptual information in the electrophysiological representational space.

#### 2.1.3. The Concept2Brain architecture: the cVAE and cross-domain neural networks

Finally, once both the cVAE (Figure 1.1) and then the cross-domain (Figure 1.2) models have been trained, the Concept2Brain model makes use of both to generate new EEG recordings from concepts not previously presented to the human participants (Figure 1.3). Specifically, the workflow of the model starts by feeding one image or text into the CLIP model (Figure 1.3 left), which calculates one 512 multidimensional vector mapping the concept representation. Next, the cross-domain network translates this conceptual representation for one participant into the electrophysiological latent space(Figure 1.3 middle). Then, the decoder module of the cVAE takes this 256 μ vector information and the one-hot 88 vector indicating the subject identification to reconstruct the EEG signal (Figure 1.3 right). Following this workflow, the Concept2Brain model can encode any picture or text semantic and emotional content into a representation of brain activity.

A deeper comprehension of the deep neural design and parameters of the model can be obtained through the open-source script accessible from https://github.com/ASantosMayo/Concept2Brain.

### 2.2. Training the Concept2Brain model

In the following section, we describe (1) the empirical data derived from human EEG recordings used to (2) fit the cVAE model to learn the electrophysiological properties of data. Then, (3) the cross-domain model is trained based on the CLIP output and the cVAE latent space.

#### 2.2.1. EEG datasets

The Concept2Brain model aims to reproduce the human neural response evoked by the visual perception of pictures. To achieve this, two EEG datasets from studies employing similar passive viewing tasks were utilized. In both studies, participants viewed a wide range of naturalistic photographs that can be categorized into three content types: pleasant, neutral, and unpleasant.

**Figure 2:**
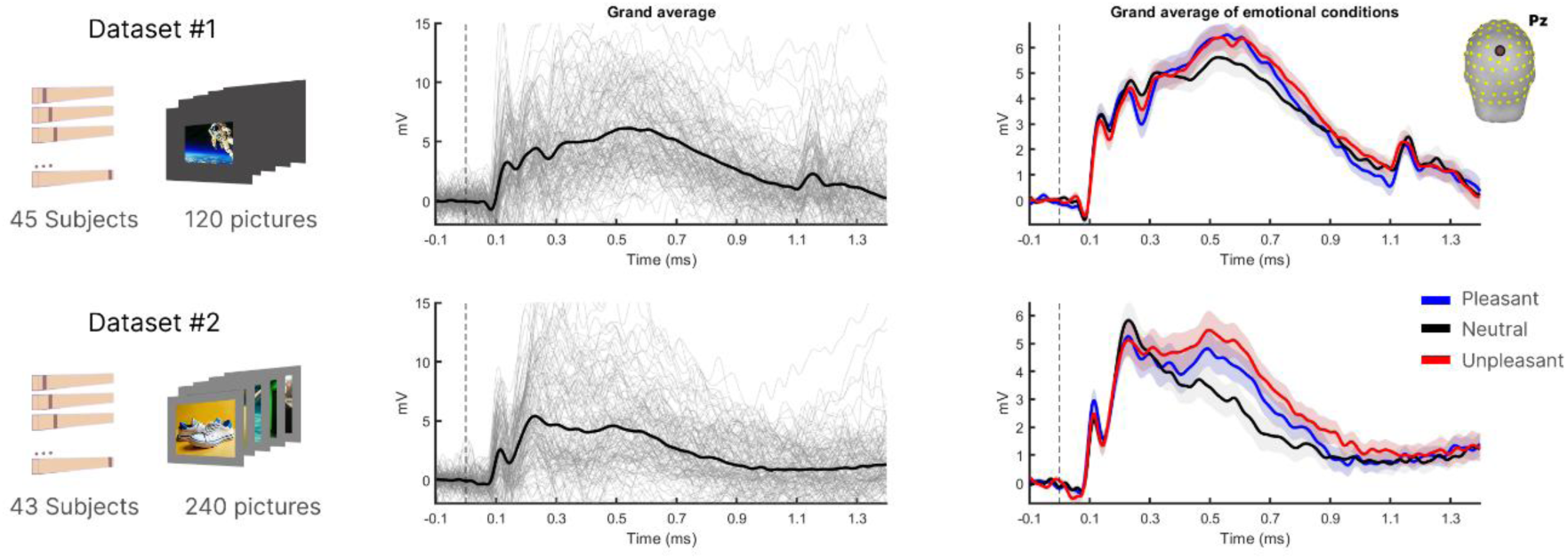
Empirical EEG datasets used for training the Concep2Brain model. Differences between both datasets illustrate differences in the presentation mode (left column): Dataset #1 involved displaying a smaller picture on a dark background (top) and Dataset #2 involved displaying a larger picture on a gray background (bottom). The middle column shows the grand average of picture-evoked ERPs, while the right column shows the averaged evoked responses by affective category (blue, red and black colors refer to pleasant, neutral, unpleasant conditions, respectively).

Dataset 1 (Figure 1 top) represents the EEG recordings of 45 subjects watching a total of 120 pictures (40 pleasant, 40 neutral, 40 unpleasant). In this experiment (Dataset #1), pictures were shown as a rectangular shape (spanning 9° of visual angle horizontally) at the center of a computer screen over a black background. Similarly, dataset 2 (Figure 1 bottom) presents the EEG recordings of 43 subjects, where a total of 240 images (80 pleasant, 80 neutral, 80 unpleasant) were presented. For Dataset 2, a gray (50% luminance) background was presented throughout the experiment and pictures were also shown the center of the screen, but spanning 15° of visual angle horizontally. Affective pictures from both studies were taken from the International Affective Picture System (IAPS)^31^ and other sources, including prior published work from the University of Georgia^32^. Each study featured a separate set of pictures, resulting in a total of 360 unique pictures across the datasets. The use of two datasets with different presentation parameters (background and foreground color, edges, and visual degrees) was intended to facilitate generalization of conceptual picture content over specific visual features.

EEG acquisition was carried out using a 128-channel HydroCel net with Ag-AgCl electrodes (impedances below 60 kΩ), connected to an EGI Net Amps 300 amplifier (sampling rate of 500 Hz). The vertex electrode (Cz) served as reference electrode. Data was preprocessed offline, by filtering using a Butterworth filter with filter orders of 3 and 9, and cut-off frequencies of 0.2 and 25 Hz. Then, EEG data was further processed by segmenting trials using 0.1 s pre-event and 1.4 s post-event, leaving trials of 1.5 s (equal to 750 time points). The Statistical Control of Artifacts in Dense Array EEG/MEG Studies (SCADS^33^) was used for artifact rejection. This method utilized the compound of the median absolute voltage, standard deviation, and max derivative of voltage for automatic pre-processing the EEG data to reject bad trials showing signal artifacts, thus ensuring the good quality of electrophysiological recordings. Data were then combined into a compound data quality (QC) index, wherein we identified channels in which the QC index was above the median of the distribution by more than 2.5 standard deviations. Those channels were then interpolated using spherical spline functions^33^. Finally, a Z-score transformation was calculated for data before incorporating it into the cVAE (see below) to facilitate smoother convergence during training. A total of 13,640 artifact-free EEG signal trials derived from 88 participants (across both datasets) were used to train the model.

#### 2.2.2. Training of cVAE model to abstract electrophysiological features

The cVAE architecture was designed using the Python-based library Pytorch^34^ (version 2.4.0). It consists of one encoder, decoder, and bottleneck blocks. Both encoder and decoder blocks are formed by 3 convolutional layers (kernel size 3), presenting a Rectified Linear Unit (RELU) activation function, commonly used to produce sparsity in the network and abstraction of non-linear information from data^35^. Afterwards, the convolutional output from the encoder layers is flattened, resulting in a 204,544-dimensional vector presenting the spatio-temporal properties of the EEG signal. To include the subject information (conditioned element of the cVAE), this vector is concatenated with another vector of size 88 containing zeros and a one in the index of the subject (one-hot vector).

All this information was condensed by two dense fully connected layers, reducing this concatenated size to two 256-dimensional vectors (μ and σ). Thus, EEG information is compressed and forced to follow a Gaussian-shape distribution where μ has values from N(0,1) and a complementary standard deviation is generated as the σ vector. This enforcement can be achieved using the “reparametrization trick”^16^, an algorithm that samples input data into this Gaussian distribution (see below). Next, the 256-dimensional μ is concatenated again with the subject vector of size 88. Mirroring the encoder, this information is converted to a 204,544-dimensional vector through two fully connected layers and finally is input into 3 convolutional layers to generate the 129 channels by 750 timepoints matrix (i.e., the synthetic EEG data).

For training the neural network, the reconstruction loss must be calculated. It represents the difference between the actual data (the real EEG signal) and the output of the neural network (the synthetic EEG signal). Here, the Mean Squared Error (MSE) was employed since both data are continuous and it is a simple and standard loss measurement. However, this reconstruction loss individually cannot ensure the correct learning of the μ and σ values, i.e. the VAE functioning. To achieve this Gaussian-shape distribution of the μ values, the Kullback-Leibler (KL)^25^ divergence was estimated to facilitate the “reparametrization trick” using the following equation:

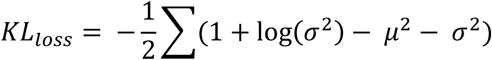

Essentially, this loss index indicates how μ (the posterior distribution) deviates from the normal distribution (the prior distribution). Therefore, to force the neural network to approximate the μ vector to follow the Gaussian distribution, the KL loss is summed to the reconstruction loss to obtain both outcomes. Nevertheless, the simple addition of the KL loss at the beginning of the training hinders the neural network from deeply exploring data to find the best reconstruction path. Therefore, the “annealing” strategy^36^ is commonly used, where the KL loss is gradually included throughout the training epochs, thus leaving the training to obtain the best reconstruction indices before starting to force the Gaussian distribution of the latent space. Here, we added an increasing KL loss across the training phase using the following sigmoid function:

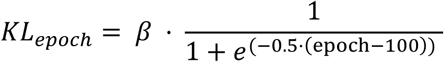

where β corresponds to 1⋅10^-5^. This parameter is needed to scale the KL loss with respect to the reconstruction loss and prevents the “posterior collapse”^25^ effect. When it occurs, μ values approach 0, resulting in a skewed distribution instead of a Gaussian-like distribution and attenuating the generative capabilities of the model.

The training phase (Adam optimizer, learning rate 1⋅10^-6^) of the cVAE was performed with a random selection of 80% of the EEG data (10,912 trials, batch size 64). A total of 1300 epochs were needed to fully train the model and obtain the electrophysiological representation of 10,912 EEG trials (randomly selected 80% of total data). A multi-GPU machine containing 3 NVIDIA RTX 4070 SUPER Ti (16 Gb vRAM) was employed. Due to model weight, encoder, bottleneck and decoder components of the cVAE were allocated in each GPU and training dataset driven for each component and GPU.

#### 2.2.3. Inference of the conceptual space and training of the cross-domain network

To create the cross-domain model, we need a latent vector of both conceptual and electrophysiological representations. In a first step, the CLIP model was employed to infer a 512-sized embedding for each of the 360 affective pictures viewed by the participants in the two EEG experiments. Second, the 10,912 EEG trials were forwarded into the cVAE model, extracting the 256-size vector μ. Hence, we have a training data set consisting of 10,912 inputs of 512 + 88 values (conceptual information) derived from the pictures and an output of 256 values.

The cross-domain neural network is composed of 3 fully-connected dense layers, presenting an exponential linear unit (ELU)^37^ activation function in the first two. Again, an Adam optimizer^38^ (learning rate1⋅10^-3^) and mean squared error (MSE) loss served to train the model. The training phase was completed within 10 epochs, when the validation loss indicated that it had reached the best model parameter estimation, while ensuring generalization (i.e., avoiding overfitting).

### 2.3. Evaluating the Concept2Brain model

Once both cVAE and cross-domain networks are tuned, the Concept2Brain model capitalizes on the CLIP abstraction of conceptual information, the cross-domain translation into an electrophysiological representation, and a decoder component of the cVAE to generate new EEG signals based on pictures or texts (Figure 1.3). To measure its performance and capabilities, two main analyses were carried out:

First, data recovery was calculated by inputting the same pictures into the Concept2Brain model as were used in the empirical studies that provided the training data. Thus, a large synthetic dataset was created, consisting of 88 pseudo-subjects and 360 pictures composed summing a total of 31,680 ERP trials. Then, an average was calculated for each subject for trials belonging to pleasant, neutral, and unpleasant conditions. A Pearson’s correlation in each channel and time point was estimated to measure the relationship between empirical and synthetic averages.

Second, we evaluated the generalization capabilities of the Concept2Brain model by employing written texts instead of pictures. To this end, 40 pleasant, 40 neutral, and 40 unpleasant short texts describing affective pictures were fed to the model for each subject. As a result, we obtained a synthetic EEG dataset of 10,560 trials showcasing electrophysiological differences within them.

Python scripts for Concept2Brain modeling, training and evaluation, along with MATLAB code for results visualization are available from: https://github.com/ASantosMayo/Concept2Brain. Trained Pytorch-based models and EEG dataset are accessible via https://osf.io/7dpgm. To enhance the visualization and interpretability of the figures in the present work, a moving average filter with a window length of 40 ms was applied to the time series depicted in this report.

### 2.4. The Concept2Brain cloud platform

The present project provides a new tool for generating open, flexible, and validated synthetic EEG datasets for the neuroscience community. We deployed our AI architecture in a cloud computing platform, allowing users to easily access the tool via the World Wide Web: https://concept2brain.brainbodybehavior.science. The web platform allows users to upload their own pictures or introduce written texts and select any number of synthetic participants at the front end (Figure 1.4 left). Note that the original participants cannot be reconstructed because of random shuffling and because of trial averaging, which obscures any fingerprint inherent in the raw EEG. Then, this information is sent to the backend server where the Pytorch model, i.e. the Concept2Brain model, generates the synthetic event-related potential (ERP) signal belonging to this input (Figure 1.4 middle). These new data are saved as a MATLAB file, a commonly used format in neuroscience research. Finally, these data are made available for download and may be visualized in a user-friendly viewer of data consisting of an interactive 3D brain and topographic graphs describing the timeseries of the 129 EEG channels across the trial. The platform allows to generate a dataset belonging only to all of the 88 synthetic participants or only one participant.

The platform was developed using React^39^, a JavaScript framework. The backend server was set using Django rest framework^40^, a python-based library used to provide the application programming interface (API) that handle user requests. Functions from the Pytorch library^34^ were utilized along with the above-detailed models to generate new EEG datasets; the MNE library^41^ was used for web-based visualization of EEG topographic results.

## 3. Results

### 3.1. Model performance and data reconstruction

First, we evaluated the model’s data recovery by measuring its performance in reproducing the EEG signal based on the same picture content. To this end, the same images used in the training data (datasets 1 and 2) were fed to the Concept2Brain model. Then, the average electrophysiological response from each one of the 88 participants was calculated for pleasant, neutral, and unpleasant content. Figure 3 shows Pearson’s correlation coefficients (r) of real and synthetic data.

**Figure 3:**
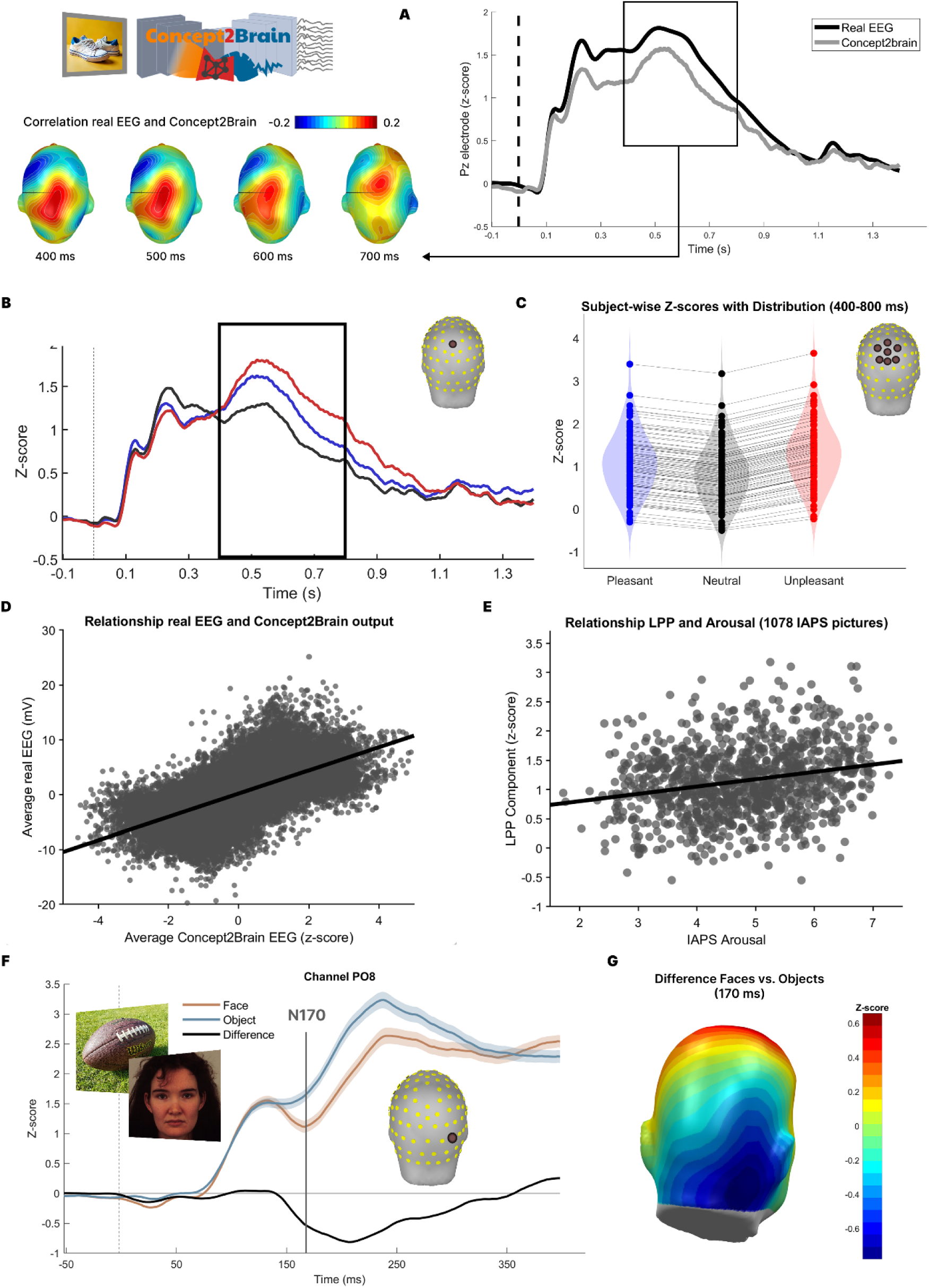
Data reconstruction performance of the Concept2Brain model. A: On the right, the plot shows in black the time series obtained from the real EEG signal (average of both datasets), and in gray the time series generated by the Concept2Brain model, both obtained from the Pz electrode. On the bottom left, topographic description of Pearson’s correlation during the 400 ms to 800 ms time window, typically associated with the LPP component. B: Time series from the Pz electrode of the synthetic EEG signal, averaged across all subjects and divided into pleasant (blue), neutral (black), and unpleasant (red) conditions. The black square shows the time range of the Late Positive Potential component (400 ms to 800 ms window), a robust measure of emotional picture content. C: Violin plot showing voltage (z-score) activity for each condition in the cluster located at the central parietal region during the above-mentioned LPP time range. All 88 subjects show a typical “V-pattern,” where emotional content (both pleasant and unpleasant) prompts greater LPP than does neutral content. D: Linear relation between real and synthetic data for each time point for the 88 subjects. E: Pearson’s correlation coefficient between normative arousal ratings collected in a standardization sample and the synthetic LPP component for 1,078 IAPS pictures. The LPP component was scored by averaging the synthetic voltage at electrode Pz across a time segment from 400 ms to 800 ms post-onset and across the 88 subjects. F: Time series of the PO8 electrode, which maximally reflects the face-sensitive ERP. Synthetic EEG signals were obtained by inputting 60 pictures of faces (light brown), 60 pictures of objects (light blue), and computing the difference between face-evoked and object-evoked waveforms (black). G: Topographic distribution of the differences between faces vs. objects at a latency of 170 ms (N170). Note that the Concept2Brain architecture exactly reproduces the canonical time course and topography of face-specific ERPs during face perception.

Results from the data recovery analysis indicate a high correlation between empirical and synthetic signals. First, the ERP response of the real EEG data and the Concept2Brain output show similar time courses and waveforms (Figure 3 A). Common ERP components such as P100^42,43^, N200^44^, P300^45^ and LPP^46,47^ are readily observed in the synthetic data, mirroring the waveform morphology of the empirical data. Next, we examined the data recovery in space and time. The highest correlations between real and synthetic signals were found in the time window from 400 ms to 800 ms post-onset of the picture (Pearson’s correlation r = 0.640, p-value < 0.01; Figure 3 E). Critically, the reconstruction accuracy shows a parietal topography during this time range. Thus, both latency and topography of the best-reconstructed synthetic data match the Late Positive Potential (LPP) component of the ERP—the most established ERP metric of emotion perception^46–48^. The Concept2Brain model effectively uses the electrophysiological information provided in the LPP spatio-temporal data segment to accurately reproduce an EEG/ERP signal corresponding to the semantic-emotional meaning of the stimulus for the observer.

Similarly to the original datasets 1 and 2, we divided the artificially generated signals into pleasant, neutral, and unpleasant categories based on the normative affective ratings of the pictures. Figure 3B, bottom, shows the difference between emotional content in the same pictures used for training the model. Crucially, waveforms of the different categories show higher neural evoked responses for the emotional pictures (pleasant and unpleasant) compared to the neutral pictures. These differences are found overall during the 400 ms to 800 ms time window, resembling those originally found in both empirical EEG recording datasets (Figure 1 right). In addition, at the individual subject level (Figure 3C), we observe, for each synthetic participant, the common “V” shaped difference of pleasant, neutral and unpleasant conditions^46,47,49^ where emotional pictures prompt greater LPP responses compared to neutral pictures. Taken together, data recovery analyses suggest that the Concept2Brain model not only is able to generate biologically realistic synthetic ERPs but also recreates late amplitude changes prototypically associated with manipulations of semantic and emotional content.

Next, we generated synthetic EEG activity of 88 subjects in response to all the standardized emotional (IAPS) pictures that were not used in the training phase of Concept2Brain. As a result, we obtained 1078 electrophysiological responses to a wide range of pictures. An extensive literature has demonstrated a robust positive linear relationship between the amplitude of the LPP component of the ERP evoked by a given picture and the self-reported emotional arousal by an observer viewing the same picture. In line with this prediction, the LPP component predicted by Concept2Brain, averaged across all synthetic participants showed a statistically significant correlation of r = 0.23 (p < 0.0001) with the normative arousal rating of 1078 pictures taken from the IAPS data base (Figure 3E).

Another validation test of the Concept2Brain’s generalizability used a canonical face-specific ERP signal referred to as the N170. It appears as a strong negativity in the ERP waveform at right-posterior electrodes, 170 ms after stimulus onset^50–52^. The N170 is substantially more pronounced (more negative) when observers view faces compared to other visual stimuli ^53,54^. To test the ability of the Concept2Brain model to reproduce this widely used effect, 60 pictures showing neutral faces (30 female, frontal and slightly oriented to camera) and 60 pictures showing common objects were selected from the KDEF^55^ and THINGS^56^ datasets, respectively, and submitted to Concept2Brain. Figure 3F shows the time course of the average Concept2Brain output across 88 subjects at electrode PO8 (right-posterior) during the first 300 ms after stimulus onset. The prototypical N170 time course and topography are both accurately reproduced, showing more negativity when viewing faces compared to the object condition. Figure 3E depicts the topographic distribution of the difference between both conditions, highlighting the involvement of right posterior-lateral areas in the N170 component, matching the topography reported in the literature^50,54^.

### 3.2. Electrophysiological response based on emotional text

Next, we applied the Concept2Brain model to text-based inputs. Leveraging the multimodal capabilities of CLIP, one of the key features of this model is its ability to extract conceptual latent representations from both images and texts. Therefore, we took this approach a step further by replicating an entire experiment, feeding the model with a total of 120 text stimuli and generating simulated ERP time series for each of the 88 virtual participants. Figure 4 illustrates the EEG signal reconstruction for all text stimuli across all participants, as well as individual reconstructions for the first three participants.

**Figure 4:**
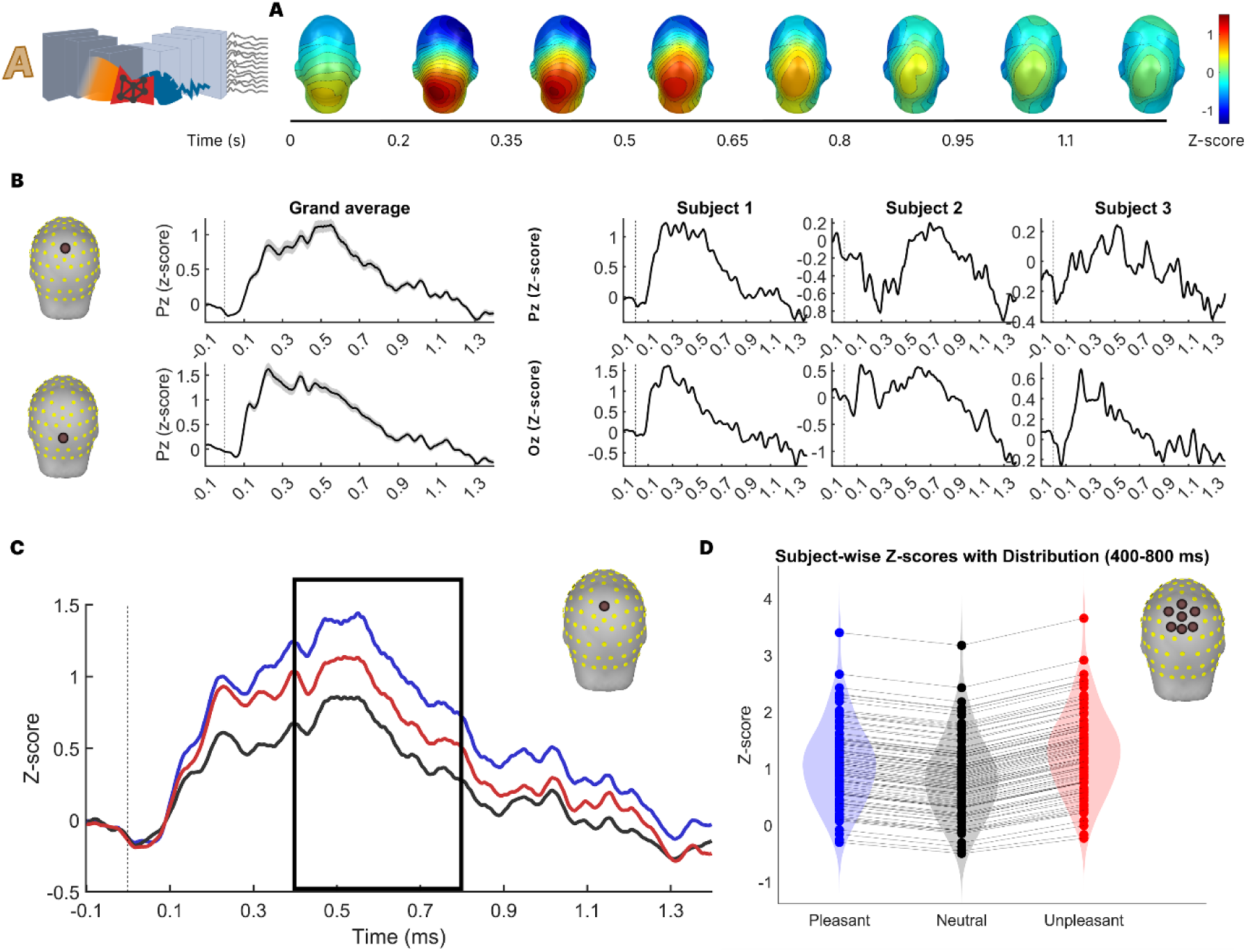
Concept2Brain generated visual ERP response to affective written texts input. **A**: Topographical distribution of the grand average ERP. **B**: ERP time series of the synthetic data for the grand average and three first subjects obtained at the Pz electrode (top row) and Oz electrode (bottom row). **C**: Time series of Pz electrode from the synthetic EEG signal averaged from all subjects and divided into pleasant (blue), neutral (black), and unpleasant (red) conditions. Black square shows the latency of the LPP component (400 ms to 800 ms window) where literature typically find differences across emotional content. **D**: Violin plot showing voltage (z-score) activity for each condition in the cluster located at the central parietal region during the above-mentioned LPP-related temporal window. All subjects (88) show the same “V-pattern” where emotional content is linked to higher activity.

First, we inspected the differences between channel waveforms. In Figure 4 A the grand average of the artificial visual ERP is shown. It clearly shows the involvement of occipital regions during the first 600 ms after the hypothetical presentation of the text-based picture. During the final part of the trial (1 s to 1.4 s), activity in the occipital sensors decreases and frontal lobes takes more interaction, suggesting post-perception higher-order processing.

To explore this further, Figure 4 B shows the simulated ERP at electrodes Pz and Oz. Distinct positive and negative going components are evident, in line with empirical data. For example, the simulated ERP at the mid-occipital Oz electrode shows deflections consistent with early visuocortical responses to stimulus onset, whereas the simulated data at the mid-parietal Pz electrode contain the late positive potential component, modulated by emotional content. Also consistent with empirical data, the ERP waveforms show pronounced differences in across participants. For illustration, three virtual participants (Figure 4 B right) show strong differences in ERP morphology—a well-established consequence of interindividual differences in brain anatomy, signal-to-noise ratio, and electrophysiological variability^57–59^.

Next, we inspected the sensitivity of the ERP activity simulated by the Concept2Brain model to the emotional content of the stimulus. Comparing ERPs in response to pictures varying in content is a widely adopted usage of ERP studies with complex, naturalistic stimuli ^60,61^. To this end, we divided the trials into pleasant, neutral and unpleasant categories based on the emotional content of the text input. Figure 4C shows the ERP at the mid-parietal electrode Pz. As expected, activity during the early time segment (from 0 ms to 200 ms) was similar between pleasant, neutral, and unpleasant categories, because this portion of the brain signal is dominated by processes related to perceptual analysis, with little sensitivity or specificity relative to emotional content. However, after 300 ms, condition differences became more evident. In accordance with empirical findings ^46,47^ (see also Figure 1), emotionally engaging cues (i.e., the pleasant and unpleasant conditions) prompt greater ERP amplitude than emotionally less engaging content (i.e., the neutral condition). Specifically, these results are observed in the parietal area during 400 ms to 800 ms (Figure 4D). Interestingly, all virtual participants, without exception, show the typically observed V-shaped pattern, with heightened LPP amplitudes for emotional stimuli^49^, with differences mainly affecting the overall level and topography of this component. Taken together, the results indicate that the Concept2Brain model robustly reproduces typical findings in EEG/ERP research.

### 3.3. ERP generation: Web implementation

In a next step, we tested the cloud computing web interface of Concet2Brain, by synthesizing the ERP to individual concepts in individual virtual participants. In an empirical study, would require the use of dozens of repeated trials using the same picture or text, to obtain a clean average of the ERP waveform. However, the Concept2Brain model allows us to estimate the averaged ERP associated with this concept in one single step for one virtual participant, because it was trained on averaged EEG activity, i.e. ERPs. As an example, we tested three concepts related to unpleasant (alligator), neutral (desk) and pleasant (beach) emotional content. These texts were fed into the model and the synthetic ERP responses are shown in Figure 5 (A and B).

**Figure 5:**
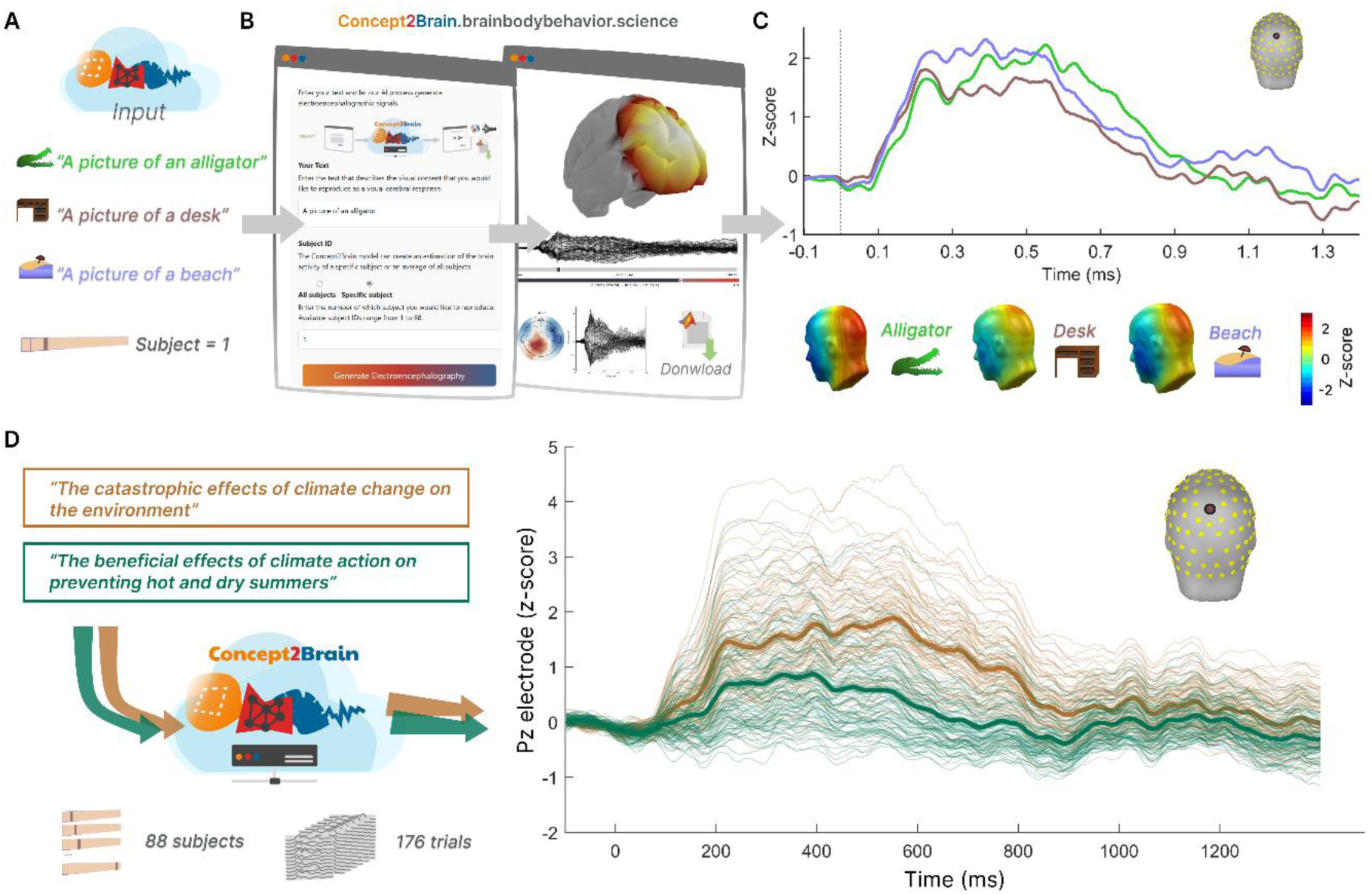
Workflow for generating individual or full-sample datasets for pictures or written texts using the cloud-computing platform of Concept2Brain. A) In the first example, simple written texts were entered into the model such as “a picture of an alligator”. In this initial example, only data for the first virtual participant were generated. B) The web platform offers an interface for uploading a picture or a written text input. Once the information is submitted, the platform generates the synthetic ERP signal. Then, the resulting data is displayed in an interactive 3D brain model across the time series along with a 2D projection, viewed from the top. The generated ERP data are readily downloaded in the MATLAB .mat format, used widely for EEG/ERP post-processing. C) These data are exemplified by the ERP waveform at electrode Pz for virtual participant number 1, showing synthetic ERPs in response to text indicating pictures of an alligator (green), a desk (brown), and a beach (blue). In the bottom row, the topography of such generated ERPs is shown for the time window 400 ms to 800 ms, illustrating how neutral content such as a desk prompts lower late ERP responses than emotionally engaging content. D) As a second example, the Concept2Brain model can also represent how 88 virtual participants respond to content related to more complex issues, difficult to implement in an empirical study. This dataset reflects two written texts describing complex content of interest and thus consists of a total of 2*88 =176 trials.

As stated above, emotional content prompts greater ERP amplitudes during the LPP time window^46,49^ (see Figure 5 C). The model produces a participant-level ERP of the that reproduces this common finding. Using the Concept2Brain platform, such differences can be examined at the level of individual prompts and individual participants, without the need of large datasets. The platform can be accessed at concept2brain.brainbodybehavior.science and generated data can be downloaded in the MATLAB file format (.mat), a widely used platform in EEG/ERP research^62^.

Finally, for illustration, we offer an example of how the platform may aid in producing synthetic data for researchers interested in complex semantic content. Such content is often difficult to study empirically, requiring extensive stimulus curation and control, while prompting substantial inter-individual differences. In Figure 5 D we illustrate the simple usage of the Concept2Brain approach for simulating brain responses to two statements about climate change. The model was used to simulate the neural response of 88 virtual participants. The synthetic data indicate heightened brain responses to statements that emphasize strong negative consequences rather than statements mentioning mitigation strategies (Figure 5D, right panel). consistent with empirical studies into well-established effects such as negative affective biases.

## 4. Discussion

Electroencephalography is the most used tool in human brain science. Its sensitivity to brain electric activity in real time makes this technique valuable for studying human behavior in a wide range of applications, from clinical assessment and treatment monitoring to economics, educational technology, entertainment, as well as the psychological and brain sciences. Here, we introduce the Concept2Brain model, a computational architecture for converting information contained in a text or image into synthetic ERP waveforms—representative brain electric responses to events.

The architecture consists of a cross-domain cVAE algorithm connecting conceptual information with brain responses. It was trained on more than 10,000 trials with 360 different pictures, representing a wide range of content. Data reconstruction analysis showed high correlation between the real ERP signal and the synthetic signal produced for the same pictures. The highest correspondence was found in central parietal regions between 400 ms to 800 ms, where the topography and latency matched the LPP component, a brain response modulated by a range of attentional and emotional processes when viewing visual stimuli^46,47,49^. Taken together, data reconstruction analysis suggests that our cross-domain cVAE architecture accurately captures the conceptual and emotional content of the pictures and generates reliable synthetic data that reproduce typical modulations of the ERP waveform by factors such as attention, motivation, and emotion, for each of the 88 subjects.

The Concept2Brain model demonstrates promising generalization abilities through its capacity to reproduce electrophysiological patterns that are well established in the literature. For example, when generating EEG activity from 1,078 standardized affective pictures, distinct from those used to train the model, the resulting LPP component showed a robust correlation with the normative arousal ratings for each picture. A second validation involved the perception of facial stimuli. Unlike objects, faces activate the fusiform face area, a brain region known for its role in face processing^53,54^. Concept2Brain accurately reproduced the characteristic topography, latency, and condition sensitivity of the N170 negative ERP deflection^50,52^, indicating that the model can learn how different input categories are reflected in EEG responses. It is critical to note that the LPP and N170 components observed in the model’s output were never explicitly programmed or introduced during training; they were instead learned implicitly from the data. Its generalization ability likely stems from the extended and comprehensive training of the cVAE model and its favorable properties. These results highlight the potential of deep learning methods to abstract subtle and complex features of electrophysiological dynamics from EEG data.

Leveraging CLIP for encoding text into an embedding space shared with pictures, we also used the Concept2Brain model to generate new ERP waveforms from written texts as inputs. Synthetic waveforms showed different electrophysiological responses for each participant, indicating inter-individual differences between virtual participants^57–59^, yet still revealing the typical ERP waveform components that are established in the literature. This finding included an accurate reproduction of the well-established modulation of the emotion-sensitive LPP component^46,49^ when manipulating emotional content of the input.

Currently, the architecture relies on the CLIP architecture from OpenAI^26^. Notably, this model allows researchers to work with cross-modal inputs, enabling them to employ both pictures and text. This architecture is suitable when images are not available, or content cannot be graphically represented. However, with continual advances in AI^27,28,63^, CLIP may not represent the most up-to-date interpreter model. The Concept2Brain methodology does not depend on this specific module, leaving open the option to use newer, more capable or more domain-specific encoders in future versions of the model.

One limitation of the present model is the limited concept space, which is based on a sample of 360 pictures. Thus, the Concept2Brain model represents not a final product, but an evolving approach to connecting state-of-the-art AI models with neuroimaging outcomes^4,64^, allowing further adaptations of this methodology with new architectures, larger datasets, or specific tasks.

Another recent advancement in signal processing is the autoencoder ^15,18,65^, which is at the core of the Concept2Brain model. In our case, the cVAE captures the electrophysiological features of the data and compresses them into 256 Gaussian-shaped variables. This electrophysiological latent space can represent the variability in the brain data, modeling a pure representation of the EEG/ERP signal obtained from brain recordings. In figure 6 we show the electrophysiological latent representation of 10,560 affective written texts obtained from the 88 subjects (see Results section 3.2).

**Figure 6:**
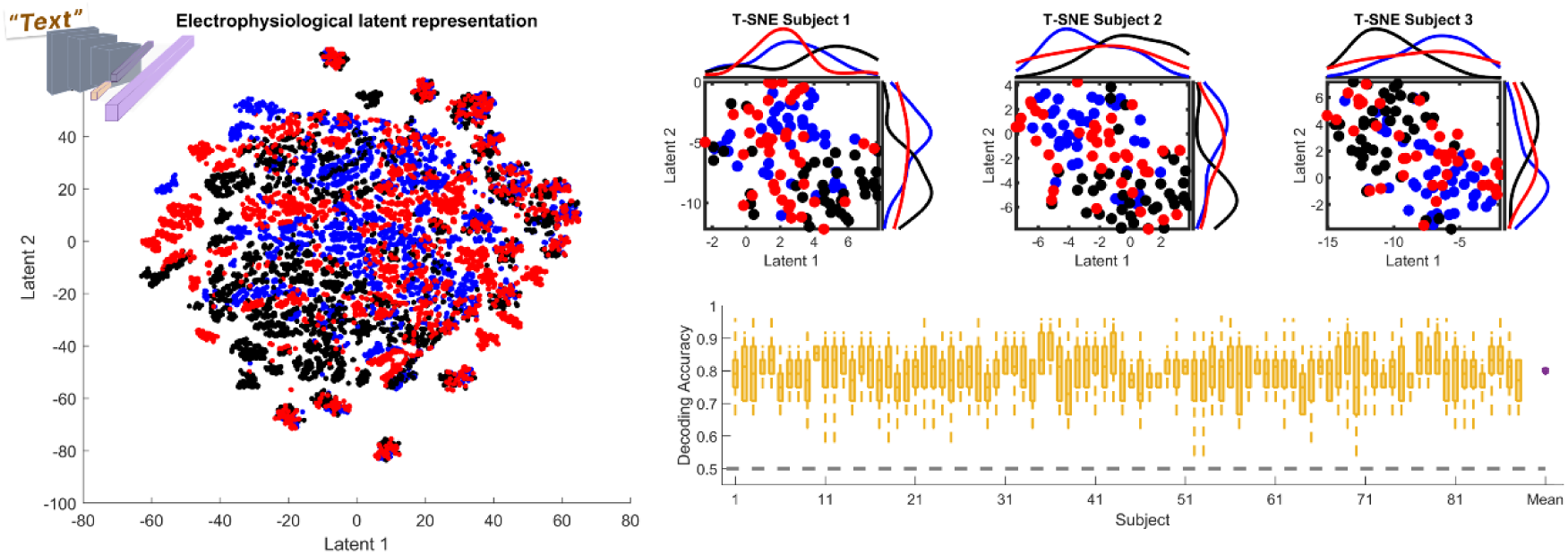
Electrophysiological latent space for affective texts. Left: Scatter plot of the electrophysiological latent space from all affective written texts projected into two dimensions, illustrated using the T-SNE method. Blue, black and red colors represent pleasant, neutral and unpleasant affective texts. Right top: Electrophysiological latent space for all written texts in the first 3 subjects. Plots show how emotional content (pleasant and unpleasant) can be distinguished from neutral content. Right bottom: A three-class support vector machine classifier was applied to the electrophysiological latent space for each synthetic participant, demonstrating high discrimination between the three affective text categories for all synthetic participants.

On the left side of Figure 6, all trials from all subjects are presented, indicating the capability of the autoencoder to reveal the idiosyncratic neural response patterns of each human brain^57,58^. Crucially, when we plot the trials obtained from each subject (top right), their electrophysiological differences for each emotion category appear. As a result of the autoencoder refinement of electrophysiological abstraction, trials showing diverse affective content can be easily decoded using support vector machine classification (SVM; Figure 6 right bottom). Taken together, this non-linear capability of deep learning shows as a promising approach for capturing complex relationships in neuroimaging data, leading to new tools for reducing its complexity and enhancing data analysis.

The Concept2Brain model represents an open-access tool aiming to offer the estimation of brain activity to many potential users encompassing a wide range of scientific and economic fields. Therefore, we deployed Concept2Brain as a cloud-computing service (concept2brain.brainbodybehavior.science) where users can utilize the model without any installation requirement. The web-based service enhances accessibility and maximizes software- and hardware independence along with making sure that users benefit from future updates of the model.

Potential uses of the Concept2Brain model span many disciplines and fields and include academic, clinical, and societal questions. For example, in clinical studies, this model may be used to generate typical brain responses for initial comparisons with a patient group. In the field of economics, the tool may contribute to modelling consumer behavior by illustrating predicted brain responses to marketing-related content^2^. In basic science contexts, the generation of synthetic EEG datasets will allow the estimation of variability and effect sizes of ERP data^57^. Finally, the absence of ecologically valid, naturalistic, or semantically complex stimuli in experimental contexts often hinders the translation of electrophysiological findings into real-life applications. The Concept2Brain model provides an avenue for addressing this challenge, generating synthetic brain waves in response to any meaningful pictorial or text input^59^. Overall, the proposed resource, with its potential to be further developed and expanded, is poised to facilitate reproducible and open simulations of brain responses to a wide range of stimuli. Thus, it enables the synthesization of explorative and reference data, as well as the initial testing of novel hypotheses, for users interested in brain responses and their modulation by semantic information.

## Acknowledgements

The authors would like to thank collaborators from the Laboratory of Brain, Body, and Behavior (LB^3^), the J. Crayton Pruitt Family Department of Biomedical Engineering at the University of Florida and the Center for Cognitive and Computational Neuroscience (C^3^N) for fruitful discussions and helpful feedback on the project. This research was supported by a grant from the National Institute of Health (R01MH125615) and by a grant from the National Science Foundation (2318984).

